# *MAXTEMP*: A method to maximize precision of the temporal method for estimating *N_e_* in genetic monitoring programs

**DOI:** 10.1101/2024.06.04.597400

**Authors:** Robin S. Waples, Michele M. Masuda, Melanie E.F. LaCava, Amanda J. Finger

## Abstract

We introduce a new software program, *MAXTEMP*, that maximizes precision of the temporal method for estimating effective population size (*N*_*e*_) in genetic monitoring programs, which are increasingly used to systematically track changes in global biodiversity. Scientists and managers are typically most interested in *N*_*e*_ for individual generations, either to match with single-generation estimates of census size (*N*) or to evaluate consequences of specific management actions or environmental events. Systematically sampling every generation produces a time series of single-generation estimates of temporal 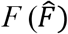, which can then be used to estimate *N*_*e*_; however, these estimates have relatively low precision because each reflects just a single episode of genetic drift. Systematic sampling also produces an array of multigenerational temporal estimates that collectively contain a great deal of information about genetic drift that, however, can be difficult to interpret. Here we show how additional information contained in multigenerational temporal estimates can be leveraged to increase precision of 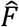 for individual generations. Using information from one additional generation before and after a target generation can reduce the standard deviation of 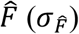 by up to 50%, which not only tightens confidence intervals around 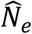 but also reduces the incidence of extreme estimates. Practical application of *MAXTEMP* is illustrated with data for a long-term genetic monitoring program for California delta smelt. A second feature of *MAXTEMP*, which allows one to estimate *N*_*e*_ in an unsampled generation using a combination of temporal and single-sample estimates of *N*_*e*_ from sampled generations, is also described and evaluated.

## 1 INTRODUCTION

Effective population size (*N*_*e*_) is one of the most important concepts in evolutionary biology but also one of the most enigmatic (Charlesworth 2009; Hare et al. 2011; Waples 2022). Because it is challenging to collect the demographic data from natural populations that is required to estimate *N*_*e*_ directly, for over half a century researchers have used genetic methods to indirectly estimate *N*_*e*_. Initially most of these estimates used the temporal method, which compares allele frequencies in samples from the same population taken at different points in time (Krimbas and Tsakas 1971; Nei and Tajima 1981; Pollak 1983; Waples 1989; Wang 2001). That changed abruptly in the late 2000s with development of two new estimators that require only a single sample: a method based on linkage disequilibrium, LD (Waples 2006, modified from Hill 1981), and a method based on the incidence of siblings (Wang 2009). Within a few years, use of single-sample estimators had far outstripped the vintage temporal method (Palstra and Fraser 2012). A recent review (Clarke et al. 2024) conducted a meta-analysis of over 4600 estimates of *N*_*e*_ from the LD method alone. This trend toward increasing interest in estimating *N*_*e*_ is likely to become even stronger in the future. Interest in regular genetic monitoring has increased in recent decades along with the realization that global ecosystems are undergoing rapid changes (Schwartz et al. 2007; De Barba et al. 2010; Foote et al. 2012; Jackson et al. 2012; Fussi et al. 2016; Van Rossum and Hardy 2022). Furthermore, *N*_*e*_ has been identified as a key metric to monitor with respect to the Convention on Biological Diversity’s post-2020 global biodiversity framework (Hoban et al. 2022; Thurfjell et al. 2022). Given the realization that global ecosystems are undergoing rapid changes, the trend toward increasing interest in estimating *N*_*e*_ is expected to continue.

These are encouraging developments, but it is unfortunate that the full potential of the temporal method is not being realized. A major feature of the temporal method is that it integrates information about effective size across all the generations spanned by the samples, such that the resulting estimate applies to the harmonic mean *N*_*e*_ over the intervening generations. Each generation of genetic drift strengthens the overall signal of effective size, so (up to a point) precision increases with elapsed time between samples. In applied conservation biology and management, however, one often wants to be able to estimate *N*_*e*_ for specific individual generations, either to match with single-generation estimates of census size (*N*) or to help evaluate consequences of specific management actions or environmental events (Kamath et al. 2015; Whiteley et al. 2015; Ruzzante et al. 2016). Single-sample estimates apply to individual generations and hence are well suited to meet these needs. If samples are taken in consecutive generations, the temporal method can also provide estimates of *N*_*e*_ that apply to specific generations. In that case, however, a single generation of genetic drift provides a relatively weak signal for estimating *N*_*e*_, so power is generally less than for single-sample estimators.

Here we show that in a genetic monitoring program that involves systematic sampling, precision of the temporal method to estimate *N*_*e*_ in specific generations can be increased by mobilizing information contained within the many multigeneration temporal estimates that also can be made. To illustrate, consider a genetic monitoring program that takes samples of progeny from generations 1, 2, …, 5 (Figure 1). Comparison of allele frequencies in samples of progeny from generations 1 and 2 provides an estimate of variance *N*_*e*_ in generation 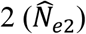, comparison of samples from generations 2 and 3 leads to 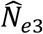, and so on. In addition, comparison of samples from generations 1 and 3 provides an estimate of the harmonic mean *N*_*e*_ in generations 2 and 3:

**Figure 1.**
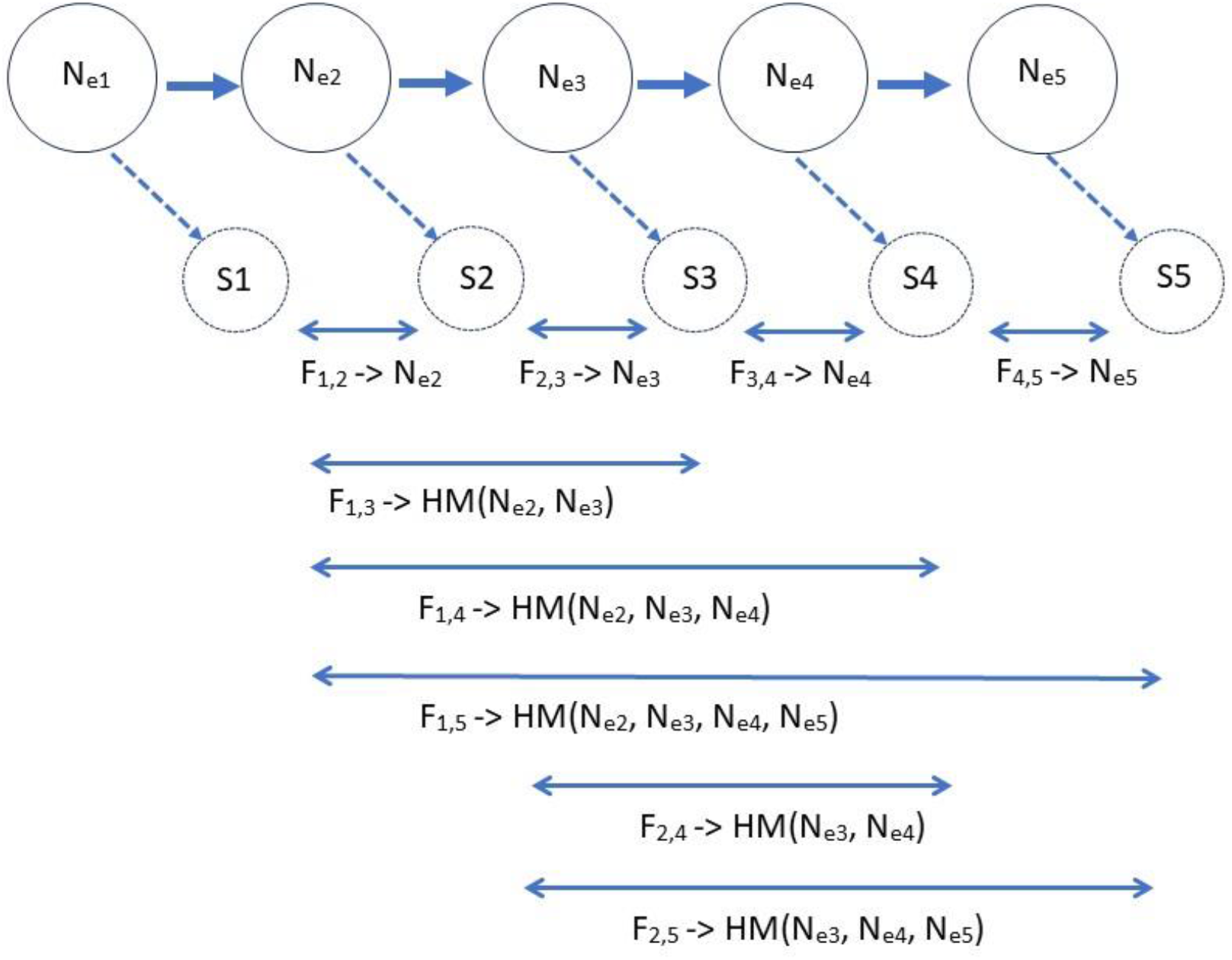
Schematic of 5 consecutive generations of samples used to estimate effective size using the temporal method. Plan II samples of *S* progeny are taken from each generation. Resulting temporal *F* values estimate single-generation *N*_*e*_ if the samples are separated by one generation. Samples separated by more than a single generation estimate harmonic mean (HM) *N*_*e*_ in the generations between samples. The long arrows at the bottom depict all temporal comparisons spanning multiple generations that include a drift signal from *N*_*e*3_.

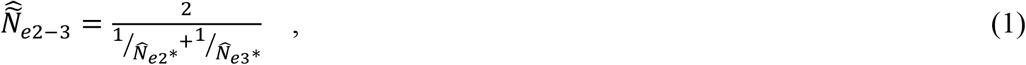

where “*” indicates an estimate that is part of a combined (multigeneration) estimate. Now we have a second estimate of *N*_*e*_ in generation 2, embedded in the right side of Equation 1. We also have a single-generation temporal estimate of *N*_*e*_ in generation 3, and if we substitute 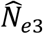 for 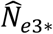 in Equation 1 and rearrange,

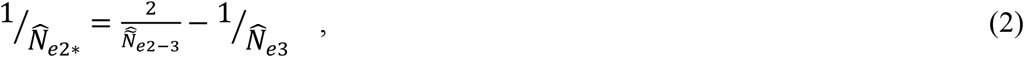

we can express the second estimate from generation 2 as a function of two temporal estimates. An improved estimate for generation 2 can then be obtained by properly weighting 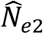 and 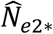, and a similar approach can be used to compute a weighted average of 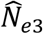 and 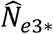. Furthermore, in theory this process can be extended to extract more information about *N*_*e*_ in generation 2 from the temporal estimates 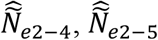, and so on, where 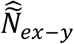 refers to the estimate of harmonic mean *N*_*e*_ for generation *x* through *y*, inclusive. With *n* consecutive samples spanning *n*-1 generations of genetic drift, there are 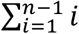 different temporal comparisons that provide information about *N*_*e*_. For example, with *n* = 10 consecutive samples, 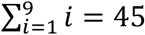 different temporal comparisons contain information relevant to *N*_*e*_ in the 9 generations spanned by the samples. However, we expect the information content to dwindle as the signal from *N*_*e*_ in the focal generation (in this example, generation 2) becomes confounded with drift signals from additional generations.

Here we use simulations to explore the feasibility of this new idea. We introduce the software *MAXTEMP*, which implements our results to maximize precision for temporal estimates of *N*_*e*_ in individual generations. We show that the percentage reduction in the standard deviation of the single-generation temporal estimator 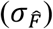 is a function of three covariates (true *N*_*e*_, and sample sizes of individuals (*S*) and loci (*L*)), and that reductions of up to about 50% can be achieved with values of *S* and *L* that are widely used. We also illustrate a second feature of *MAXTEMP*, which deals with incomplete sampling. In a long-term study there are often data gaps, which (despite the best intentions) can arise for a variety of reasons (e.g., lapse in grant support; departure of key personnel; freezer meltdown; random screw-ups; global pandemic).

Consider the above scenario but without the sample from generation 2. The temporal estimate 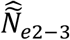, based on samples from generations 1 and 3, provides an estimate of harmonic mean *N*_*e*_ in generations 2 and 3. If a single-sample method can be used to obtain 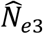 for generation 3, that single-generation estimate can be used in Equation 2 to obtain an estimate of *N*_*e*_ applicable directly to generation 2.

## 2 METHODS

### 2.1 Temporal estimation of *N*_*e*_

The temporal method for estimating *N*_*e*_ relies on the fact that the standardized variance in allele frequencies comparing two samples separated by *t* generations of genetic drift (tempF, hereafter just *F*) is a simple function of *N*_*e*_ and the sample sizes of individuals at times 0 and *t* (*S*_*0*_, *S*_*t*_) (Nei and Tajima 1981; Waples 1989):

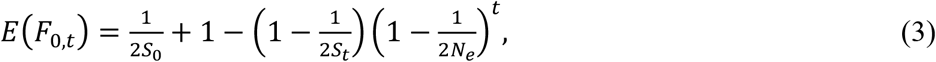

where *E*(*F*_0,*t*_) refers to the expected value of *F* calculated for samples of progeny from generations 0 and *t*. Equation 3 applies to sampling Plan II of Nei and Tajima (1981) and Waples (1989), which requires that the initial sample of individuals that are progeny of reproduction in generation 0 cannot be involved in reproduction in generation 1. If any individuals in the initial sample can contribute genes to subsequent generations, then sampling is according to Plan I and it is necessary to add a term to Equation 3 for *N* (census size at the time of the initial sample) to account for correlations of allele frequencies over time. We don’t attempt that here, but in theory it could be done subsequently.

When elapsed time is small compared to *N*_*e*_ (*t*/*N*_*e*_ << 1), *F* increases approximately linearly by magnitude 1/(2*N*_*e*_) per generation, and a good approximation is:

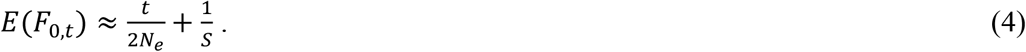

Rearrangement of Equation 4 produces an estimator of *N*_*e*_:

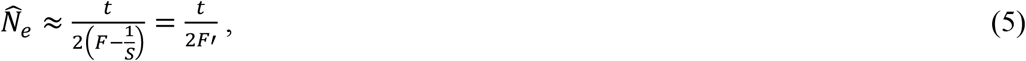

where *F*′ = *F* – 1/*S* is the raw temporal *F* adjusted for sampling a finite number of individuals. If *S* differs between the two samples, the harmonic mean of the two sample sizes should be used in the above equations.

The above equations apply to a constant *N*_*e*_ and need to be modified to allow *N*_*e*_ to vary over time. Assume now that the true *N*_*e*_ values for each generation are labeled *N*_*e1*_, *N*_*e2*_, etc.

Then Equation 3 can be modified as

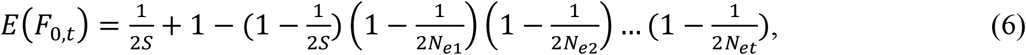

which can be approximated by

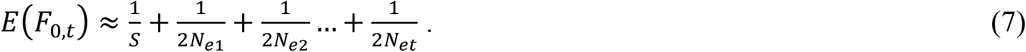

For each of the single-generation *N*_*e*_ terms in Equation 7, we can substitute the respective value for *F* spanning a single generation:

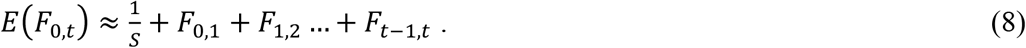

This is very useful, as the distribution of 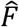 is well-known and well-behaved, whereas the distribution of 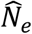 is highly skewed and can take biologically impossible (infinitely large) values. Hereafter we deal with 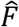 for estimation and only convert to 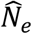 after all adjustments to 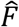 have been made.

Now focus on estimating *N*_*e*_ in a single generation, using the example in Figure 1. Let’s say we want to use all available information to estimate *N*_*e3*_. The 5 generations of samples can be used to generate a triangular matrix of 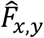 values spanning 1 to 4 generations of genetic drift (Table 1). The most direct way to estimate *N*_*e3*_ is to calculate 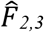 for samples of progeny from generations 2 and 3, as that estimator is not affected by a drift signal from any other generations. But there is also information about *N*_*e3*_ in other pairs of samples that span generation 3: samples 1,3, 1,4, 1,5, 2,4, and 2,5. Consider the samples from generations 1 and 3, which produce the estimator 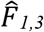. From Equation 8,

**Table 1.** Triangular matrix of temporal estimators that can be computed, given the experimental design in Figure 1, which involves sampling in 5 consecutive generations. 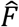 values along the diagonal can be used to estimate single-generation *N*_*e*_; off-diagonal elements provide an estimate of harmonic mean *N*_*e*_ across multiple generations. With *n* consecutive samples, there are 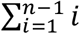 (which is 10 for this example) different temporal *F* values that provide information about genetic drift relevant to *N*_*e*_ in the *n*-1generations between samples.

| Generation of<br>Sample 1 | Generation of Sample 2 |  |  |  |
| --- | --- | --- | --- | --- |
|  | 2 | 3 | 4 | 5 |
| 1 | $\hat{F}_{1,2}$ | $\hat{F}_{1,3}$ | $\hat{F}_{1,4}$ | $\hat{F}_{1,5}$ |
| 2 | | $\hat{F}_{2,3}$ | $\hat{F}_{2,4}$ | $\hat{F}_{2,5}$ |
| 3 | | | $\hat{F}_{3,4}$ | $\hat{F}_{3,5}$ |
| 4 | | | | $\hat{F}_{4,5}$ |

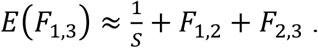

Solving for the drift signal specific to generation 3 produces another temporal estimator:

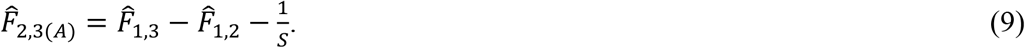

This same procedure can be used to generate a full series of estimators applicable to *N*_*e3*_:

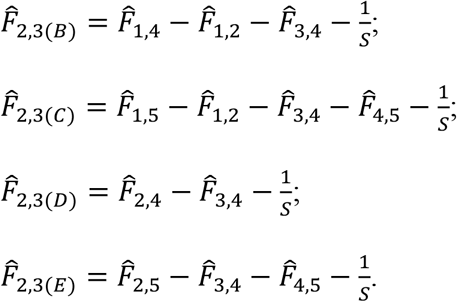

To get an overall estimate, we give each separate estimator a weight (*W*) that is the reciprocal of its variance. Then, the overall temporal *F* estimator for *N*_*e*_ in generation 3 is

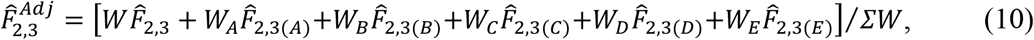

where “*Adj*” indicates an adjusted estimate that includes information from additional generations. The overall estimator of *N*_*e3*_ is then

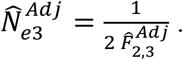

### 2.2 Precision

Temporal *F* has a distribution that is a multiple of a χ^2^ distribution (Nei and Tajima 1981; Pollak 1983), but the distribution becomes effectively normal if at least 50 independent alleles are used to compute 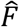. Here we consider diallelic (SNP) loci, which have one independent allele per locus, and we assume that enough loci are used that 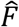 can be considered to be normally distributed. This means that confidence intervals (CIs) for 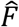 can be calculated as a simple function of the standard deviation of 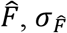. For example, the 95% CI for 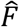 would be

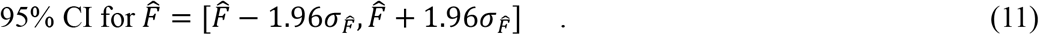

CIs for 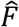 are then used (as in Equation 5) to generate CIs for 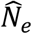.

Under pure drift, after Lewontin and Krakauer 1973, the expected variance of 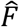 for *L* diallelic loci is 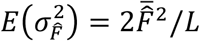, so the expected (parametric) standard deviation is 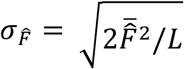. This provides a useful reference point for evaluating increases in precision obtained by using more of the information contained in multigeneration temporal estimates. Using computer simulations (described below), we empirically measured 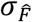 across replicates for single-generation temporal estimates and compared that with 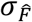 for adjusted estimates that also used information from one or more multigeneration temporal estimates that spanned the focal generation.

### 2.3 Computer simulations

All simulations were done in R (version 4.2.2; R Core Team 2022). Three estimators of temporal *F* are commonly used: 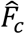 (Nei and Tajima 1981); 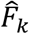 (Pollak 1983); and 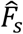 (Jorde and Ryman 2007). As a first step, we evaluated which estimator would be most suitable for use in our study. We used a Wright-Fisher model to simulate allele frequency change across many replicate 5-generation time series of constant or variable *N*_*e*_ ranging from 50 to 500. The evaluation criterion we used was root-mean-squared-error (RMSE), which reflects both bias and precision and is computed as the square root of the mean squared difference between the estimate and the true parameter. For this purpose, true *F* was taken to be 1/(2*N*_*e*_) for the respective generation. Consistently, 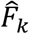 had the smallest RMSE, smaller even than was found for any of the three pairwise means of the estimators, or for the overall mean of all three estimators. Accordingly, all remaining evaluations used Pollak’s estimator. For diallelic loci, temporal *F* is the same regardless which allele is used, and 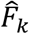 is computed as

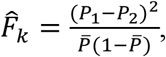

where *P*_1_ and *P*_2_ are frequencies of the focal allele at times 1 and 2 and 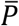 is the mean of *P*_1_ and *P*_2_. Mean 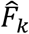 is computed as a mean across loci, with monomorphic loci (with 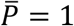 and 1 − 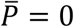) being excluded. In what follows, we drop the subscript *k* as all 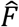 apply to Pollak’s estimator.

Subsequent analyses all simulated a series of 7 consecutive generations of Wright-Fisher reproduction, with separate (Type II) samples of progeny taken for genetic analysis from the parents in each generation. The 5 middle generations formed the Core analyses, as these generations all had at least one generation of sampling before and one generation after to draw on for additional information regarding genetic drift. Generations 1 and 7 were Edge generations, as their estimates could be improved by considering either one generation before or one generation after, but not both.

The general approach to obtaining an adjusted estimate of 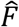 and its variance was iterative. For focal generation *X*, an adjusted estimator 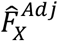 was obtained as illustrated in Equation 10. This was repeated for each generation in the time series. At this point, the whole process was repeated, as we now had improved estimates of 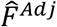 for each individual generation to use in Equation 9. The next sets of simulations were designed to answer several key questions:

1. How many generations away from the focal generation provide useful information that improves the estimates?
2. What number of iterations produces optimal results?
3. Should only the single-generation 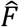 values be updated each iteration (the diagonal elements in Table 1), or should the multigeneration (off-diagonal) estimates be updated as well?
4. What is the optimal weighting scheme for multiple estimators of 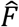 for the same generation?

Tackling Question 1 first, we picked generation 3 in the middle of the time series as the focal generation. For each replicate, we computed the single generation estimate 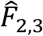, as well as the full suite of multigeneration estimates that included generation 3. From the latter, we constructed additional estimates of 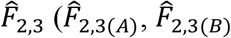, etc.) using the approach outlined in Equation 9 and subsequent material. Across all replicates, we computed 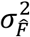 for each estimator and constructed a vector of weights (*W*) that were inverses of 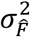 —hence theoretically optimal weighting, assuming the estimators are independent. Finally, using these weights, for every replicate we computed adjusted 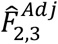 values for composite estimates that included 1, 2, 3, or 4 additional generations of data, and across all replicates we computed 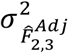 for all these different adjusted estimates. Inspection of the results showed that adding information from one additional generation before and after the focal generation reduced 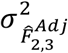, but that including data for generations farther removed did not. Accordingly, the remainder of our analyses concentrated on the focal generation, plus one generation on either side.

Questions 2-4 were considered jointly in a modified simulation scheme. *N*_*e*_ for the 5 middle (Core) generations took different sets of values ranging from 50 to 1000, and for each replicate these values were randomly scrambled to minimize effects of the order of population sizes. Regardless of the scrambled order, data were tracked separately for each true *N*_*e*_ value. The answer to Question 3 rapidly became clear: updating the off-diagonal elements consistently reduced the standard deviation of the adjusted estimates (Figure S1). For Questions 2 and 4, we used RMSE of 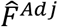 as an evaluation criterion and applied it to the same random series of true *N*_*e*_ values used for Question 3. For Question 4, previous results had shown that optimal weights for the 3 estimators of focal *F* were closely approximated by either ***W***=[0.3, 0.4, 0.3] or ***W***=[0.25, 0.5, 0.25], where the middle value applies to the single-generation 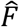 and the other two weights apply to estimates using one additional generation of data. Results of simulations evaluating these two weighting schemes with 2-4 iterations of updating estimates showed that optimal performance was found for 3 iterations using the [0.25, 0.5, 0.25] weighting scheme (Figure 2), so these values were used in subsequent analyses. For the Edgegeneration estimates, which were based on only two estimators—the single-generation 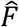, and another that included data from one additional generation either before or after—optimal weighting schemes were close to ***W***=[0.4, 0.6] or ***W***=[0.6, 0.4], with the larger weight applying to the single-generation 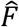.

**Figure 2.**
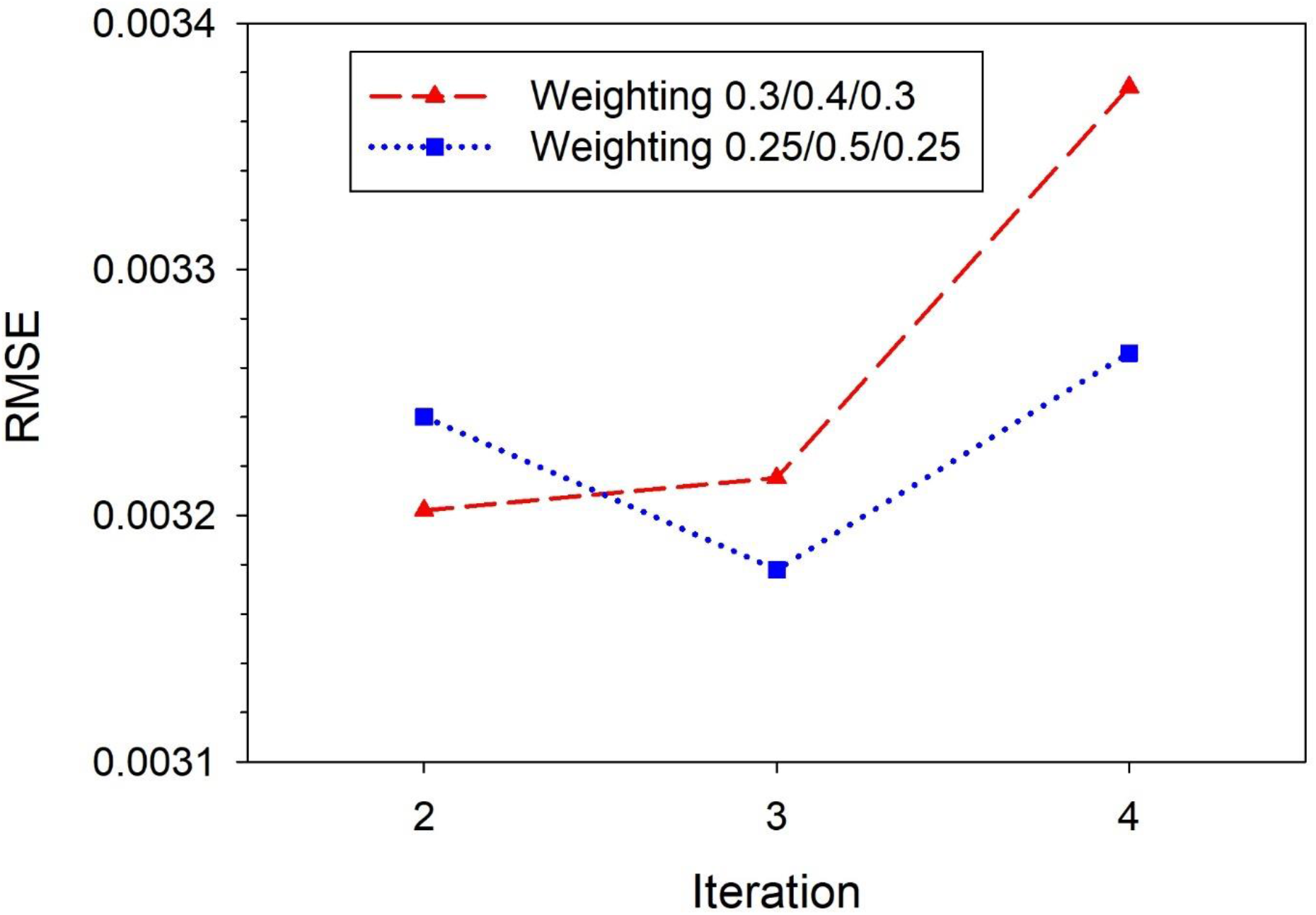
Summary of simulation results evaluating the optimal weighting scheme and the optimal number of iterations for updating 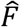. RMSE is root-mean-squared error of 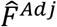, averaged across the same sets of *N*_*e*_ values shown in Figure S1. The middle weight is for the initial single-generation 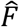 for the focal generation; the left and right weights are for estimates of *F* in the focal generation derived from samples that also reflected one additional generation of genetic drift before or after the focal generation, respectively.

**Figure 3.**
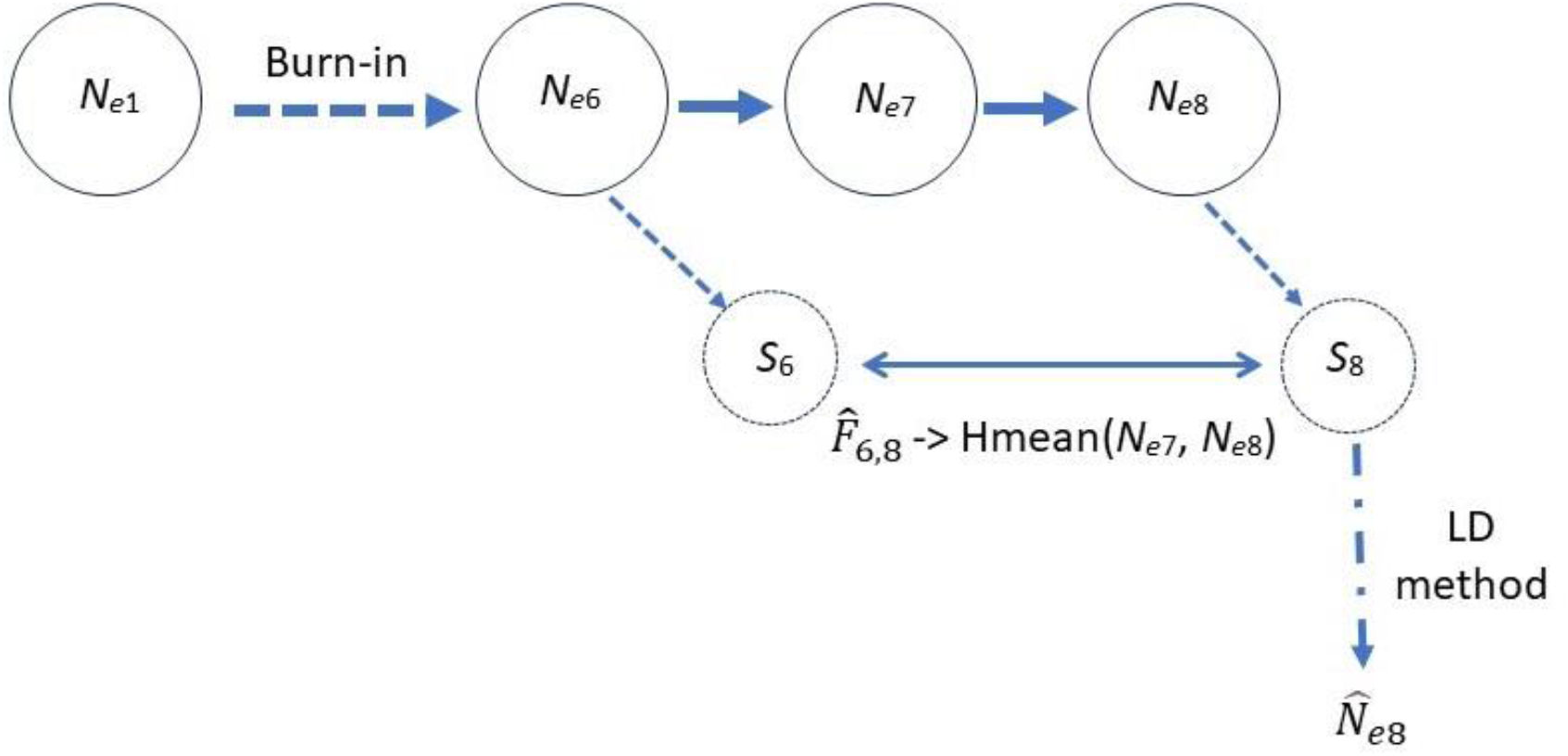
Schematic of the scenario used to model a missing sample. Samples of offspring are taken from generations 6 and 8 but not 7. Temporal *F* comparing allele frequencies in the two samples 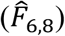 provides an estimate of the harmonic mean variance *N*_*e*_ in generations 7 and 8. If a single-sample method (in this case the LD method) can be used to estimate *N*_*e*8_, the “mystery” *N*_*e*_ in unsampled generation 7 can be estimated using Equation 2.

With these preliminary evaluations out of the way, we were ready to evaluate performance of *MAXTEMP* for a range of parameter values for key covariates of interest to researchers, which (in addition to *N*_*e*_) include sample sizes of individuals (*S*) and loci (*L*, assumed here to be diallelic SNPs). The simulation framework was similar to the above, with one difference involving the range of *N*_*e*_ values. In the preliminary evaluations, we considered up to a 20-fold range of true *N*_*e*_ within each simulation to approximate values found in different populations that might be of interest to researchers. At this point, however, we want to focus on maximizing usefulness of *MAXTEMP* for application to genetic monitoring programs for a single population, and that magnitude of variation in abundance would be excessive in most cases.

Accordingly, for each simulation scenario we picked a single Target*N*_*e*_ and modeled variation above and below that target. True *N*_*e*_ in generations 1 and 7 were set to Target*N*_*e*_. Within the 5 Core generations, three were set to Target*N*_*e*_, one was set to 2*Target*N*_*e*_, and one was set to Target*N*_*e*_/2, with the order being randomly scrambled each iteration. This scheme modeled a population whose effective size varied 4-fold over a short time span of 5 generations. We simulated all possible combinations of the following covariate values: *S* = [25, 50, 100, 200]; *L* = [100, 600, 2000]; *N*_*e*_ = [50, 75, 100, 150, 200, 300, 400, 500, 650, 800, 1000]. For each of the 132 parameter combinations, we ran 1000 replicates and recorded data for all generations with true *N*_*e*_ = Target*N*_*e*_.

Results of these simulations were used to model the relationship between the response variable 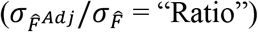 and the 3 covariates. Ratios for 3 of the 5 Core generations that equaled the Target*N*_*e*_ were treated as replicates in the modeling. Based on exploratory plots of the simulated data, we regressed log-transformed Ratio on log-transformed covariates *N*_*e*_, *S*, and *L*, including a quadratic term for log-transformed *N*_*e*_:

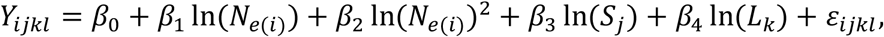

where *Y*_*ijkl*_ = natural log-transformed Ratio 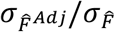 and 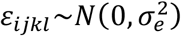.

We also ran simulations to illustrate the second capability of *MAXTEMP*: compensating for missing data. The LD method (Waples and Do 2008) was used to generate single-sample estimates of *N*_*e*_. We used a Target*N*_*e*_ of 100 and ran 6 generations of burn-in to allow levels of LD to reach a dynamic equilibrium, followed by 2 more generations of drift. Multilocus genotypes were tracked for 1000 unlinked loci. In the Constant scenario, Generations 7 and 8 also simulated Target*N*_*e*_, whereas in the Change scenarios, Generations 7 and 8 simulated either *N*_*e*_ = 125 and 75, respectively (Change/Up), or *N*_*e*_ = 75 and 125 (Change/Down). Samples of *S* = 25, 50, or 100 offspring were taken of progeny from generations 6 and 8 but not 7. The samples from generations 6 and 8 were used to compute 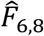, which when adjusted for sample size and used in Equation 5 with *t* = 2, was expected to produce 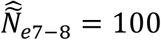 for the Constant scenario and 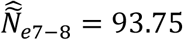 (the harmonic mean of 75 and 125) for the two Change scenarios. The expected value of estimated *N*_*e*_ for the LD sample from generation 8 was either 75 (Change/Up) or 125 (Change/Down), while the true (mystery) *N*_*e*_ for unsampled generation 7 was the reverse (125 for Change/Up and 75 for Change/Down). Following Equation 2, Mystery *N*_*e*_ for generation 7 was estimated as

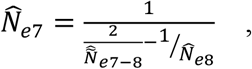

where 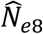 was the LD estimate and 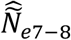 was derived from 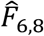.

### 2.4 Empirical example

We illustrate practical application of the new method using empirical data for a long-term genetic monitoring program for delta smelt, *Hypomesus transpacificus*, which is endemic to the San Francisco Estuary in California, USA. The delta smelt is a useful system for testing *MAXTEMP* because the species has a nearly annual life cycle (Moyle et al. 1992) and represents a single genetic population (Fisch et al. 2011). In addition, due to the complexity of and human impact on the San Francisco Estuary, a variety of systematic, long-term monitoring surveys have been conducted in the area, some for decades (Tempel et al. 2021). Methodologies for these surveys are largely unchanged over time, and survey crews record all species captured and take samples as needed, providing a rich archive of DNA from delta smelt that can be used for genetic analysis. In particular, estimating *N*_*e*_ has been of great interest to managers, especially as the wild population nears extinction and a program of experimental releases from the conservation hatchery has begun (USFWS 2020; 2022).

For this example, we used previously generated RAD-sequencing data for wild delta smelt obtained in state and federal trawls from 2013 to 2020 (SRA BioProject PRJNA1112857; https://www.ncbi.nlm.nih.gov/sra). For more details on sample sourcing, sequencing and data processing, see Supporting Information. We used 1114 SNP loci for samples of 50 individuals each for the years 2013-2020, inclusive (Table S1). Samples were assigned to cohorts for *N*_*e*_ estimation based on survey timing and gear used. Delta smelt typically spawn in February–May (Moyle et al. 1992). For example, samples assigned to the 2013 cohort are offspring from spawning events in 2013. The full matrix of temporal 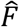 values using Pollak’s estimator were computed in NeEstimator (Do et al. 2014). For each year (= each generation), we used *MAXTEMP* to compute adjusted 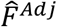 values, which were used to provide improved estimates of *N*_*e*_ each generation. Based on predictions from our linear model, given the specific covariate values applicable to each year, we generated improved CIs for each of these 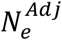 point estimates.

## 3 RESULTS

### 3.1 Adjusted temporal estimates

The best-fit model used natural log to transform 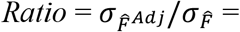 the ratio of the standard deviation of the adjusted 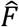 to the standard deviation of the initial, single-generation 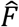 and all covariates. All covariates were highly significant, and the model explained 92.1% of the variance in the data. The estimated prediction equation for *Ratio* was

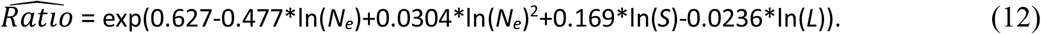

The response variable, 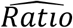, allows one to estimate the percent reduction in 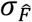 that is possible from utilizing more of the data:

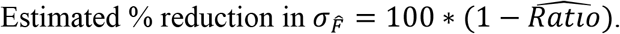

As shown in Figure 4, for *N*_*e*_<100 the percent reduction in 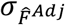 compared to 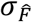 is only about 10-30%, depending on values of the other covariates, but for moderate to large 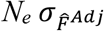 can be reduced by nearly half.

**Figure 4.**
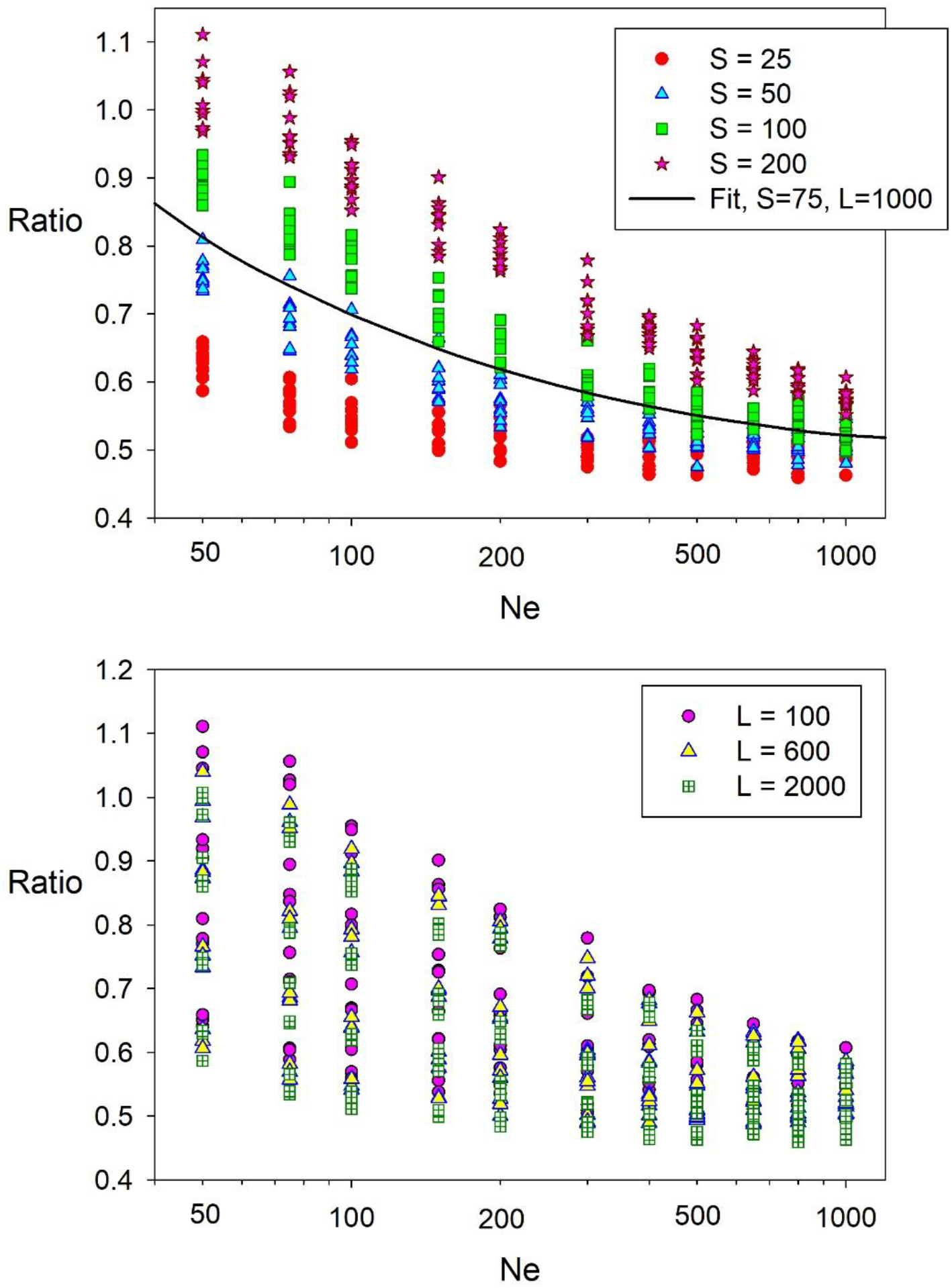
Simulation results quantifying increases in precision for 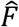. Symbols are color coded according to sample size (*S*, top) or number of loci (*L*, bottom). 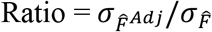 is the ratio of the standard deviation of 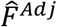 to the standard deviation of the initial, single-generation 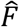. The black curve in the top panel is the predicted 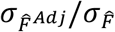 assuming *S* = 75 and *L* = 1000. Note the log scale on the X axis. These results are for estimates that can leverage information from one additional generation of genetic drift both before and after the focal generation. See Figure S2 for results for estimates that can use information from one additional generation of genetic drift either before or after the focal generation, but not both.

Hypothetical data in Figure 5 illustrate the dramatic effect the reduced variance associated with 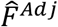 can have for CIs around 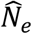. Assuming accurate point estimates for true *N*_*e*_ from 50 to 800, the parametric CIs using only the single-generation 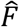 are shown on the left and the adjusted CIs that take into consideration one additional generation of data on either side of the focal generation are shown on the right. Relative improvement of the CIs increases with *N*_*e*_ and is particularly noticeable for *N*_*e*_ = 200. For *N*_*e*_ = 400, the adjusted CI has an upper bound of just under 2000, while the upper bound to the parametric CI is infinity. For *N*_*e*_ = 800, the upper bound of both CIs is infinity, but the adjusted CI has a considerably higher lower bound (329 vs 205).

**Figure 5.**
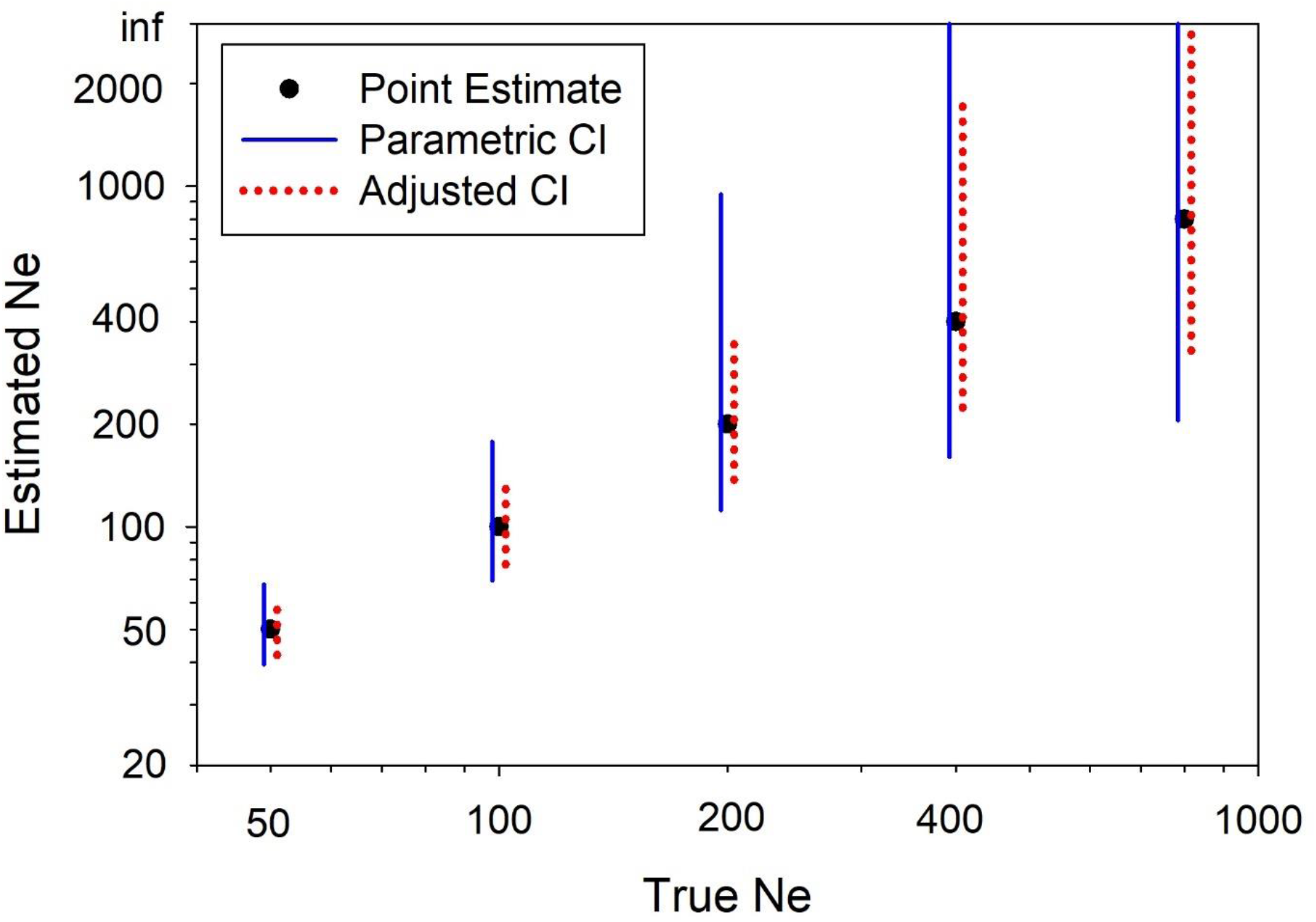
Point estimates and confidence intervals for hypothetical scenarios with *S* = 50, *L* = 1000, and true *N*_*e*_ = 50-800. 95% CIs are arranged around point estimates (black circles) assumed to agree with true *N*_*e*_. CIs on the left side of point estimates (solid blue lines) are based on parametric expectations for 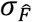, and CIs on the right (red dotted lines) are based on modeled 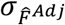. Note the log scale on both axes.

These results assume that at least one generation of additional data are available both before and after the focal generation. When the focal generation is at either the start or end of a genetic monitoring program, the potential benefits are reduced. In that scenario, the best fit model was

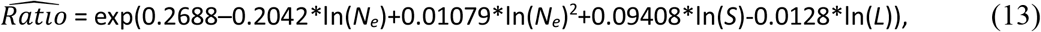

with an adjusted *R*^2^ of = 0.849. With moderate to large *N*_*e*_, the maximum percent reduction in 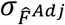 is about 30% rather than ∼50% when the focal generation is in the middle of the time series (Figure S2).

### 3.2 Empirical example

Increased precision for temporal estimates provided by *MAXTEMP* is nicely illustrated in the delta smelt example (Figure 6). After adjustment, CIs were narrower for 6 of the 7 generations. The only exception was 2014, for which only one extra datapoint was available to improve the estimate; the adjusted point estimate also increased considerably, which (all else being equal) leads to wider CIs. The direct benefits of *MAXTEMP* are most easily seen for generations 2015 and 2018. In 2015, the adjusted point estimate is slightly higher, but nevertheless the upper bound of the CI is finite, whereas it extended to infinity in the original data. In 2018, the initial and adjusted point estimates are nearly the same, but the width of the adjusted 95% CI is much narrower (148-480 vs 110-1424). The new results also have direct conservation relevance. The lower bounds of all adjusted CIs were higher, which is good news from a conservation perspective. For the initial estimates, lower bounds of 6 of the 7 initial CIs dipped below 200, and 7th (for 2019) almost did. After applying *MAXTEMP*, only 2 adjusted CIs dropped this low.

**Figure 6.**
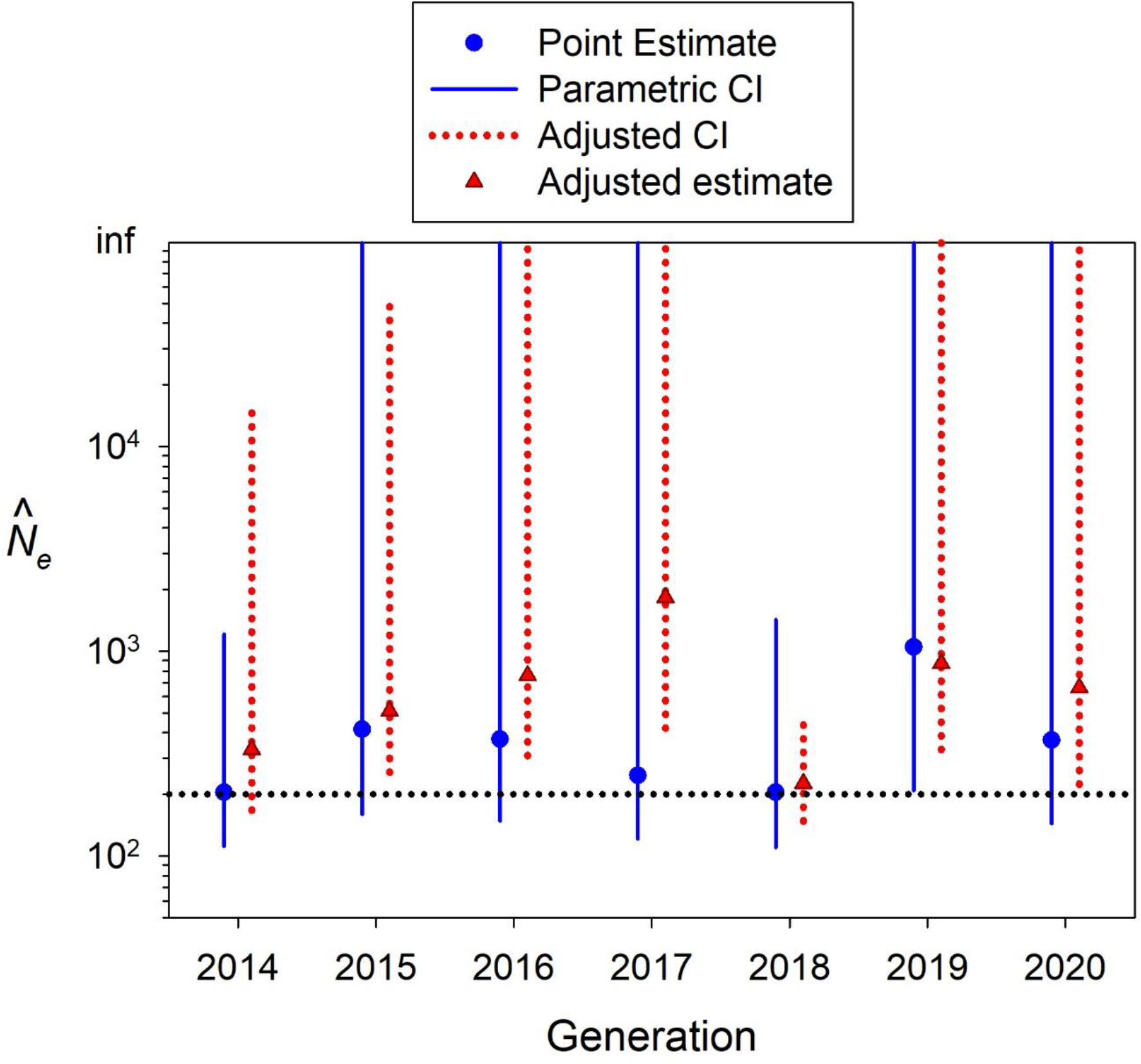
Point estimates of *N*_*e*_ and 95% confidence intervals for 7 generations of delta smelt data. Initial point estimates and CIs on the left (blue circles and solid blue lines) are based on initial 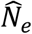 and parametric expectations for 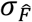, and adjusted point estimates and CIs on the right (red triangles and red dotted lines) are based on 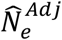 and modeled 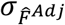 Upper bounds of some CIs extend to infinity (inf). Note the log scale on the Y axis.

### 3.3 Compensating for a missing sample

#### 3.3.1 Bias

In the Constant *N*_*e*_ scenario of the missing-sample method (Figure 7, top), the temporal-method estimate for generations 7-8 was very close to the expected value of 100, whereas the LD estimate for generation 8 was on average about 3-5% too high. With these two estimates as inputs, the estimate of Mystery *N*_*e*_ in the unsampled generation 7 was a bit low on average, from about 3% for *S* = 100 to about 10% for *S* = 25.

**Figure 7.**
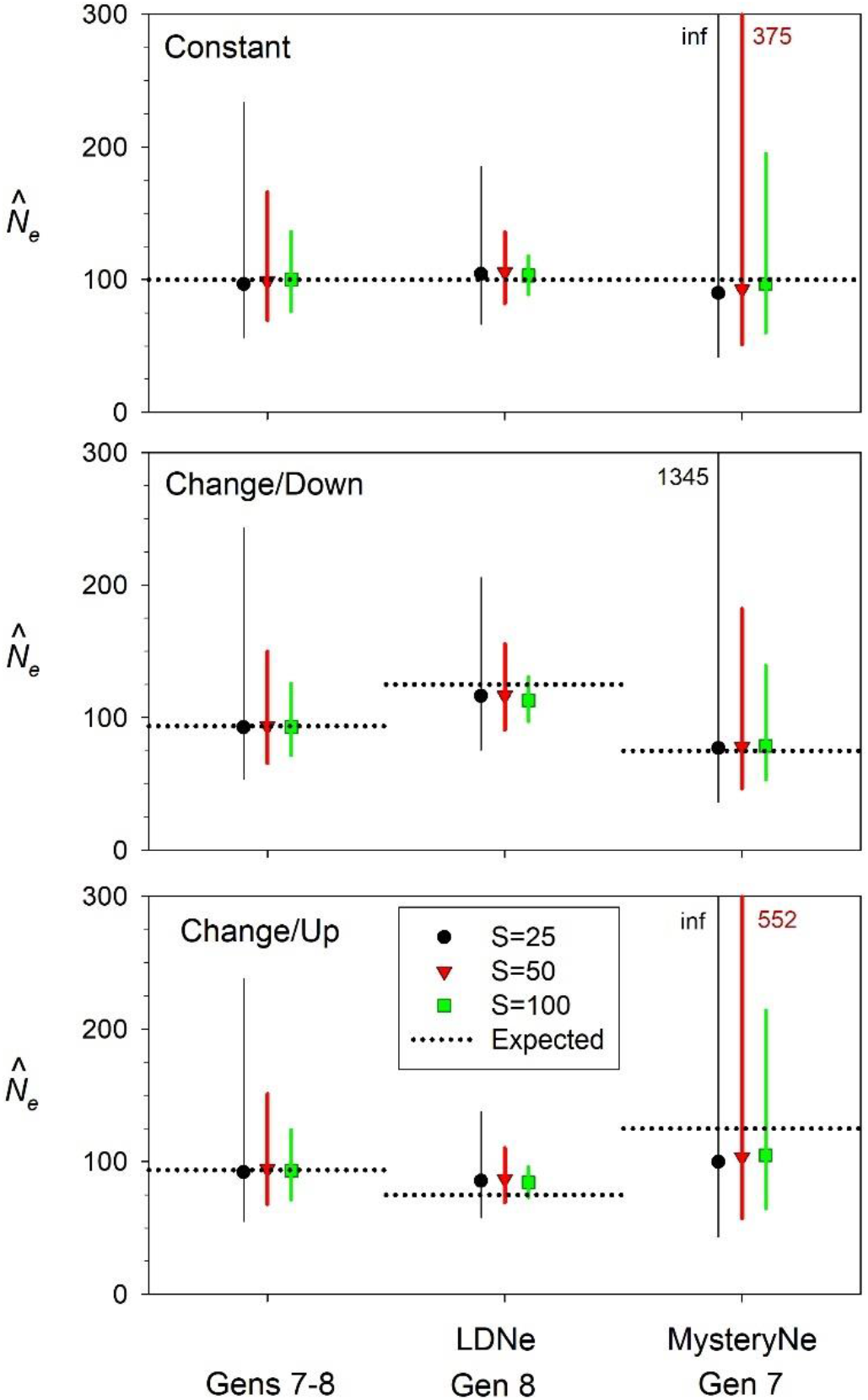
Results of performance evaluation of the method to estimate *N*_*e*_ in an unsampled generation. See Figure 3 for a schematic representation of the experimental design. True *N*_*e*_ in generations 6-8 is either [100/100/100] (top panel), [100/75/125] (middle), or [100/125/75] (bottom). Samples of *S* = 25, 50, or 100 progeny are taken from generations 6 and 8, but not 7, and genotyped for 1000 diallelic loci; symbols show resulting harmonic mean 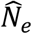 across 1000 replicates. Samples from generations 6 and 8 were used to compute 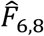 and this was used to estimate the harmonic mean *N*_*e*_ in generations 7 and 8 (left). The sample from generation 8 was used to estimate *N*_*e8*_ with the LD method (middle), and this information was used in conjunction with information from the temporal samples to estimate Mystery *N*_*e*_ in the unsampled generation 7 (right). Horizontal dotted lines represent true *N*_*e*_ for generations 7 and 8 and harmonic mean true *N*_*e*_ across both generations. Vertical colored lines for each sample size show empirical 95% confidence intervals for 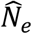; colored numbers indicate upper CI bounds that exceeded 300.

In the population-change scenarios (Figure 7, middle and bottom), the temporal-method estimate was very close to the expected 94, which is the harmonic mean of 75 and 125. In the Change/Down scenario, where *N*_*e*_ first dropped to 75 in generation 7 before climbing to 125 in generation 8, the LD estimate for generation 8 was slightly low for all sample sizes, and the estimate for the unsampled generation 7 was essentially unbiased (Figure 7, middle). With the reverse pattern (Change/Up, where *N*_*e*_ rose to 125 before declining to 75), the LD method overestimated *N*_*e*8_ by about 10-15%, and the estimate of Mystery *N*_*e*7_ was correspondingly biased downwards (Figure 7, bottom).

#### 3.3.2 Precision

Consistently, empirical CIs to 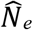 were tightest for generation 8 using the LD method and widest for the unsampled generation 7. As expected, regardless of the method, CIs were narrower for larger sample sizes. Reduced precision from small samples was particularly noticeable for Mystery *N*_*e*7_. With *S*=25, upper CI bounds were >1000 for all three scenarios and infinitely large in two of the three, whereas with *S*=100 the upper CI bounds were all <215.

## 4 DISCUSSION

Two previous studies have incorporated information from multiple samples to obtain temporal estimates of *N*_*e*_: Pollak (1983) for the standard temporal method that assumes discrete generations, and Jorde and Ryman (1995) for a modification that accounts for age structure and overlapping generations. However, both of these methods are designed to estimate a single *N*_*e*_, which is assumed to be constant over time. In contrast, *MAXTEMP* is designed to maximize information regarding *N*_*e*_ in specific generations. Major themes related to practical implementation of the new method are summarized below.

### 4.1 Adjusted temporal estimates

By leveraging additional information about genetic drift contained in multi-generation estimates of temporal *F* that are routinely generated as part of a genetic monitoring program, the standard deviation of 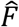 applicable to individual generations can be reduced by up to about 50% for generations that are interior to the series and by up to about 30% for the first and last generations. Proportional increases in precision are greatest when true *N*_*e*_ is large and sample sizes of individuals are small. This is good news for researchers, as those are the scenarios for which it is most difficult to obtain robust estimates of effective size. All else being equal, a reduction to 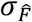 also reduces the size of confidence intervals around 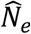. For example, with samples sizes of *S* = 50 individuals scored for *L* = 1000 SNPs and true *N*_*e*_ = 200, expected 95% confidence intervals to 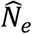 would be 112-947 (range = 835) based on 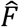 spanning a single generation but would drop to 137-367 (range = 230) after integrating information from two other multi-generation estimates (Figure 5).

In addition to narrower CIs, a second benefit to the new method is to stabilize the point estimates of 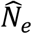, such that on average they span a narrower range. In any given empirical dataset, this general pattern might be manifest in a variety of ways. In the delta smelt example (Figure 6), all but one of the adjusted estimates were higher than the raw 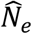. This result, combined with tighter adjusted CIs for every generation except 2014, presents a slightly more favorable picture of the species’ conservation status than did the raw data.

It should be pointed out that, regardless whether raw or adjusted estimates are used, there is a positive correlation between 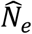 and CI width, such that (for fixed sample sizes of individuals and loci) CIs are wider when 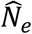 is larger. For example, for the standard temporal method using data shown in Figure 5, parametric 95% CIs are 40-68 for true *N*_*e*_ = 50 and 160 to infinity for true *N*_*e*_ = 400. This means that if adjusted point estimates of *N*_*e*_ increase using *MAXTEMP*, adjusted CIs might also be wider, even if 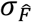 is reduced. This effect is seen for 2014 in the delta smelt example (Figure 6): the point estimate for that year increased by over 50% (from 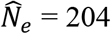 to 330) and the lower bound of the CI rose sharply as well (from 111 to 167), but the upper bound for the adjusted CI rose from 1208 to almost 15000.

If 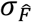 for individual generations continued to decline as more and more extra generations of data were used, long-term genetic monitoring programs could be particularly effective in increasing precision of temporal estimates of *N*_*e*_. That, however, proved not to be the case, as using more than one additional generation on either side of the focal generation did not continue to increase precision. We expect that this result reflects the non-independence of estimators using overlapping sets of the same generations. The default inverse-variance weighting scheme is optimal only if the elements being weighted are independent, but that is not the case here. We are interested in the variance of the adjusted estimator 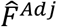, which is the weighted sum of two or more terms (call them *A* and *B*). Standard theory tells us that var(*A*+*B*) = var(*A*) + var(*B*) + 2cov(*A*,*B*). In our experimental design, the pairwise covariance terms are positive and therefore affect the optimal weighting scheme. Consider, for example, a design with samples of progeny taken from 3 consecutive generations, 1-3, leading to the single-generation estimator 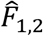 and the multi-generation estimator 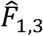. These two estimators can be used as described above to derive the adjusted estimator 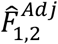, which then can be used to estimate *N*_*e*_ in generation 2. Now consider a fourth sample, from generation 4, and the new estimator 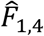, which contains information about genetic drift transitioning from generation 1 to 2, from 2 to 3, and from 3 to 4. However, 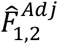 already includes 2 separate estimates of genetic drift for generations 1-2 and one estimate for generations 2-3. Furthermore, 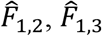 and 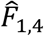 all use the same sample from generation 1 to establish initial allele frequencies. All of these factors create a large positive covariance between the drift signal contained in 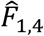 and information that is already reflected in 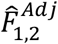. In effect, these covariance terms create a barrier that must be surmounted before a new estimator usefully can be added into the mix. In our preliminary modeling, positive covariances associated with any estimators extending more than one generation beyond the focal generation were large enough that attempting to include them in the analysis degraded rather than improved performance.

#### 4.1.1 Genomics-scale datasets

Analytical and simulation results presented here have assumed that the loci used are all unlinked and hence provide independent information about genetic drift. In reality, all loci have to be apportioned among a relatively small number of chromosomes (mean = 25 in vertebrates; Li et al. 2011), within which recombination is relatively rare. As more and more loci are used to estimate key genetic metrics like temporal *F*, precision does not increase as fast as it would if all the datapoints were independent. Waples et al (2022) quantified the effects of this lack of independence (aka pseudoreplication) on precision of mean *r*^2^ (a measure of LD) and *F*_*ST*_. The latter has distributional properties similar to temporal *F*, so the Waples et al. (2022) results for *F*_*ST*_ are relevant here. If the loci are all independent, the variance of 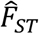 (and 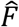) should be inversely proportional to the number of diallelic loci, *L*: 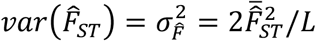. In this equation, *L* can be considered to be the degrees of freedom associated with the estimate. Waples et al. showed that the actual (effective) degrees of freedom for *F*_*ST*_ (*L*′) was a function of true *N*_*e*_, *S, L*, and genome size (number of chromosomes), and that using *L*′ rather than *L* in the above equation accurately predicted the true variance of 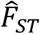. Unless *N*_*e*_ is small (<100), *L*′ is appreciably less than *L* only when more than 1000 loci are used, and if *N*_*e*_ is moderately large (∼1000) there is little pseudoreplication unless about 10^4^ or more loci are used (Figure 8). Hence, lack of independence is expected to have had little effect on the analyses reported here. However, large genomics-scale datasets can have 10^4^-10^7^ loci even for non-model species, so this issue is important to consider more generally. *MAXTEMP* incorporates results from Waples et al. (2022) that allow it to predict *L*′ and *L*′/*L* for temporal *F*, given known values of *S* and *L* and estimates of *N*_*e*_ and genome size.

**Figure 8.**
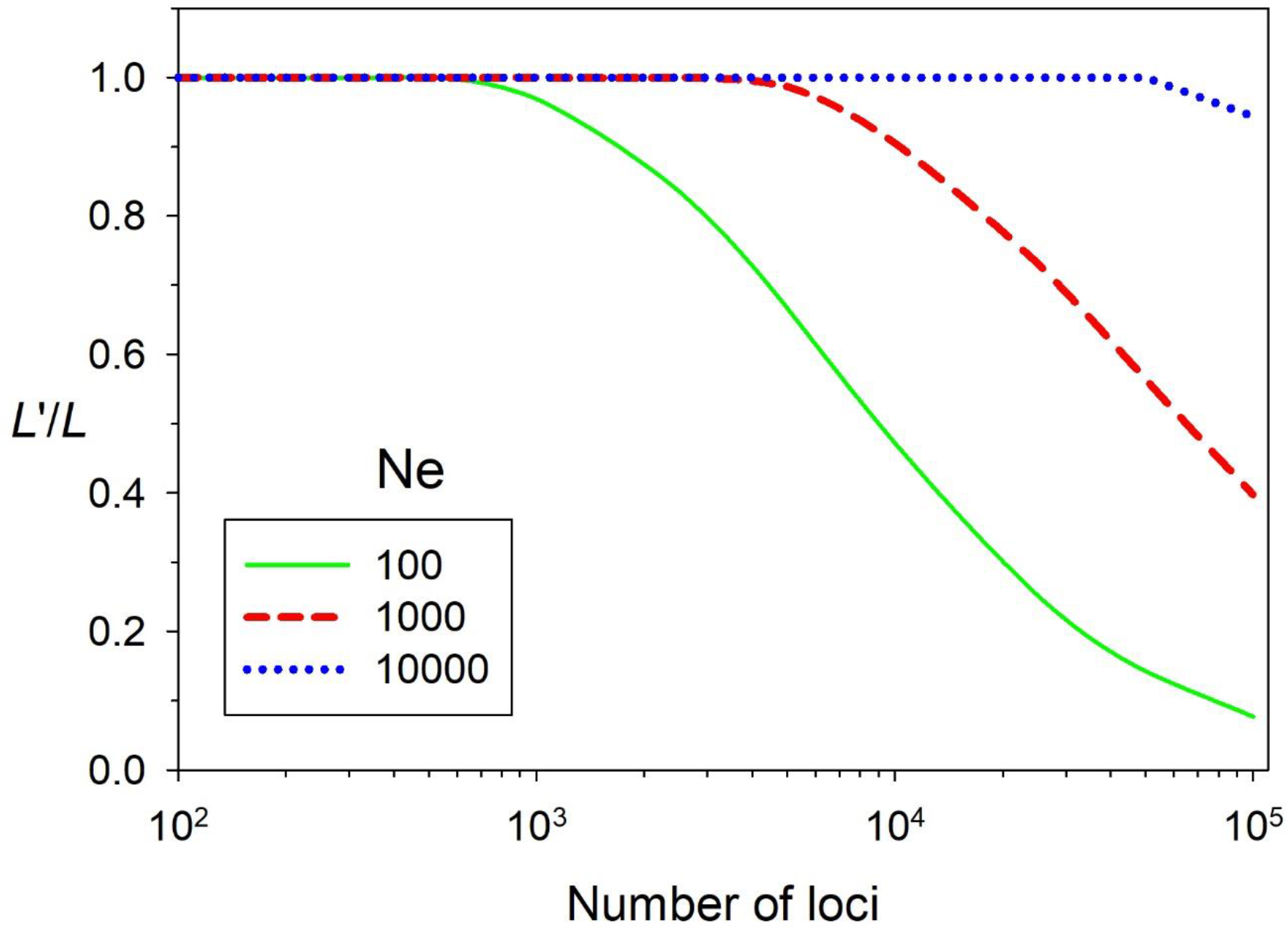
The ratio of the effective number of diallelic loci used to estimate *F*_*ST*_ (*L*′) to the actual number of loci (*L*), as a function of *N*_*e*_ and *L. L*′ is defined as the number of theoretically independent loci that would produce the same variance of mean *F*_*ST*_ as the *L* loci actually used. Put another way, the *L* linked loci have the same information content as would *L*′ totally independent loci. Plots are based on results from Waples et al. (2022), assuming the species has 20 chromosomes and samples sizes are 50 individuals. The distribution of temporal *F* is similar to that for *F*_*ST*_ so results should be applicable to analyses reported here.

### 4.2 Compensating for a missing sample

Collecting samples from natural populations is often logistically challenging, time consuming, and expensive, so occasional gaps are not uncommon in genetic monitoring programs. Here, we showed how a one-generation gap can be overcome, using a combination of temporal and single-sample estimators to obtain an estimate of *N*_*e*_ in the unsampled generation. In our model, the temporal method was used to estimate harmonic mean *N*_*e*_ in generations 7 and 8, and these estimates were largely unbiased, both for constant *N*_*e*_ and variable *N*_*e*_ (Figure 7, left). Accuracy of the estimates of mystery *N*_*e*_ in the unsampled generation 7 therefore depends on accuracy of the single-sample estimate for generation 8. With constant *N*_*e*_, the LD method tends to slightly overestimate effective size, which leads to a corresponding slight underestimate of *N*_*e*_ in generation 7 (Figure 7, top). When *N*_*e*_ changes over time, the LD method is influenced to some extent by *N*_*e*_ in previous generations. Under the Change/Down scenario (true *N*_*e*_ = 75 in generation 7 and 125 in generation 8), the LD estimate for generation 8 was biased slightly downwards by the lower *N*_*e*_ in generation 7, but the estimate of mystery *N*_*e*_ showed essentially no bias (Figure 7, middle). Under Change/Up (true *N*_*e*_ = 125/75; Figure 7 bottom), the higher *N*_*e*_ in generation 7 pushed the LD estimate for generation 8 higher, and (combined with the slight overall tendency to overestimate *N*_*e*_) this caused the overall LD estimate for generation 8 to be about 10-15% too high. This in turn caused an underestimate of *N*_*e*_ in unsampled generation 7.

General conclusions regarding bias that emerge from our results are as follows. If *N*_*e*_ is constant or nearly so, or if *N*_*e*_ is lower in the unsampled generation, the estimate of mystery *N*_*e*_ will be unbiased or nearly so, with somewhat better performance for larger sample sizes. If *N*_*e*_ is larger in the unsampled generation, the two potential biases in the LD method reinforce each other, with the result that mystery *N*_*e*_ can be underestimated to some extent. In this case, an option would be to use Wang’s (2009) sibship method to provide the single-sample estimate of *N*_*e*_. The incidence of siblings should not be affected by *N*_*e*_ in previous generations, which could lead to less bias.

It is apparent from Figure 7 that there is a cost in terms of reduced precision for having to estimate Mystery *N*_*e*_ without the benefit of a directly-relevant sample. Fortunately, this reduced precision can be overcome to some extent by taking larger samples of individuals from other generations.

### 4.3 Caveats and limitations

Several caveats are in order regarding the new methods proposed here. First, like the standard temporal method and most other genetic methods for estimating *N*_*e*_, *MAXTEMP* assumes that generations are discrete. Most species in nature are age-structured, and age structure introduces potential biases that depend, among other things, on the experimental design and the species’ life history (Waples and Yokota 2007). Second, the treatment here assumes sampling is according to Plan II, where individuals comprising the initial sample are collected before reproduction and removed so that they cannot contribute genes to subsequent generations. If the initial individuals are sampled after reproduction or non-lethally before reproduction, sampling is according to Plan I and it is necessary to add a term for *N* into Equations 3-5, where *N* is the total number of individuals subject to sampling the first sample (Nei and Tajima 1981; Waples 1989). The model could be modified to account for Plan I sampling, but that is not attempted here.

Third, the standard temporal method assumes a closed population, and estimates can be biased if immigrants with different allele frequencies enter the population between sampling events.

Effects of migration are generally relatively small over short time periods unless the migration rate is fairly high (Luikart et al. 2010; Gilbert and Whitlock 2015). Finally, the temporal method is based on the theoretical drift variance in frequency of neutral alleles, and estimates can be biased by strong directional or stabilizing selection. Researchers might consider using one of the many software programs that attempt to identify ‘outlier’ loci, which show unusually high levels of allele frequency change over time. Filtering out such loci might provide a more reliable indicator of random genetic drift.

## Supporting information

sample data

## Acknowledgments

Genetic work for delta smelt was supported through Bureau of Reclamation Grants #R10AC20089, #R15AC00030 and #R20AC00027, and State Water Contractors Agreement A19-1844. Delta smelt samples were collected by the University of California, Davis Fish Culture and Conservation Laboratory, the California Department of Fish and Wildlife, and the U.S. Fish and Wildlife Service. Delta smelt genetic data were generated by Alisha Goodbla, Emily Funk, Mary Badger, and Grace Auringer. The authors would like to thank Jim Hobbs and the Otolith Geochemistry and Fish Ecology Lab at UC Davis for assistance with assigning delta smelt samples to cohorts. The scientific results and conclusions, as well as any views or opinions expressed herein, are those of the author(s) and do not necessarily reflect those of NOAA or the Department of Commerce.

## Data Availability

Genetic data and associated metadata used for delta smelt data are deposited in the SRA (BioProject PRJNA1112857; https://www.ncbi.nlm.nih.gov/sra). A version of *MAXTEMP* that analyzes the delta smelt data is included in Supporting Information. R code to conduct the simulations and analyses in this paper will be posted on Zenodo on acceptance.

