## Supplementary material for "*MAXTEMP*: A method to maximize precision of the temporal method for estimating *N_e_* in genetic monitoring programs": Suppl Info

for

### Empirical example

*Sample acquisition*

We collected and sequenced a total of 1,152 archived delta smelt samples obtained in state and federal trawls from 2013–2020 (Table S1). Samples were grouped into generation cohorts based on collection date and life stage. Details on sample sources are in the table below and additional details on survey design can be found in Mitchell et al. (2019) and Tempel et al. (2021).

| Source | Long Name | Sampling Period | Age Group | Organization |
|---|---|---|---|---|
| EDSM | enhanced delta smelt monitoring program | fall | sub-adult | U.S. Fish & Wildlife Service |
| FCCL | UC Davis fish conservation and culture laboratory | winter/spring | adult | University of California, Davis |
| FMWT | fall midwater trawl | fall | sub-adult | California Department of Fish and Wildlife |
| CCE | Covered cod end study | fall | sub-adult | California Department of Fish and Wildlife |
| SKT | spring kodiak trawl | spring | adult | California Department of Fish and Wildlife |
| STN | summer tow net | summer | juvenile | California Department of Fish and Wildlife |

*Sequencing and data processing*

Genomic DNA was extracted using Qiagen DNeasy Blood and Tissue Kit (Qiagen, Valencia, CA) according to the manufacturer's protocol. In order to produce a large number of loci in a cost-effective manner, restriction site associated DNA (RAD) sequencing was carried out for all individuals. RAD libraries were prepared using the *Sbf1* restriction enzyme according to the 'new RAD protocol' described in Ali et al. (2016). Sample years 2013-2017 and 2018-2020 were sequenced with 100 bp and 150 bp paired-end reads, respectively, on an Illumina HiSeq 4000. We demultiplexed the sequencing data using previously described perl scripts to separate raw sequencing data into plates, then individuals (Ali et al. 2016). We trimmed 150 bp reads to 100 base pairs to have more comparable data across sample years. We then aligned the individual sequencing files to the Delta smelt reference genome (GCA_021917145.1) using bwa (Li and Durbin 2009), which resulted in sequence alignment map (SAM) files. We then further processed the SAM files using SAMTOOLS (Li et al. 2009) by sorting according to read name (samtools sort), filling in mate coordinates (samtools fixmate -m), removing duplicate reads (samtools markdup -r), and indexing the resulting files to create binary alignment map (BAM) files (samtools index) for downstream analyses.

*Principal component analysis for hybrid detection*

Delta smelt have been observed to hybridize with wakasagi smelt (*Hypomesus nipponensis*) in the San Francisco estuary (Benjamin et al. 2018). Due to the possibility of visual misidentification or technical error, we ran a principal component analysis to identify and exclude hybrid individuals. To do this, we used the program ANGSD (Korneliussen et al. 2014) to randomly sample a single read at all sites contained in at least half of the samples for each individual (angsd -doMajorMinor 1 -minMapQ 20 -minQ 20 -SNP_pval 1e-12 -GL 1 -doMaf 1 -doCov 1 - doIBS 1 -doCounts 1). This creates a 0 to 1 matrix for each individual's sampled allele at all locations in the form of a covMat file. We then calculated obtained eigenvalues using the program R (R Core Team 2013), calculated the observed variance for PC1 and PC2, and visualized the first and second principal components (PC) and removed outlier individuals.

*Data preparation for $N_e$ estimation*

In order to standardize the number of gene copies contributing to the estimation of $N_e$, we subsampled to 50 individuals from each birth year. We then called genotypes in the selected individuals using allele frequency as priors in ANGSD. SNPs meeting the following criteria were accepted: posterior probability greater than 0.85 (-postCutoff 0.85), a SNP p-value greater than 1e-6 (-SNP_pval 1e-6), found in greater than 50% of individuals (- minInd 1472), minimum mapping quality of 20 of greater (-minMap 20), minimum base quality of 20 or greater (-minQ 20), and a minimum minor allele frequency of at least 0.05 (-minMaf 0.05), and genotypes were written as numbers (-doGeno 2) in a geno file. The geno file was read into R for further filtration using the snpR package (Hemstrom and Jones 2022). Within snpR we filtered SNPs that violated Hardy-Weinberg Equilibrium (HWE=0.99), did not have read coverage in at least 75% of individuals in each year (min_ind=0.75). We also used the reference assembly to select SNPs that were not within 1Mbp of each other to reduce bias in from linkage.

*Results*

We identified and removed a total of 8 individuals identified as potential hybrids. ANGSD identified a total of 20,956 loci, which we further filtered in R to produce a final dataset containing 1,114 loci for $N_e$ estimation.  Not all loci were variable in every temporal comparison.

Table S1. Delta smelt data used for the empirical example. yr1 and yr2 are years of first and second temporal samples. gen1 and gen2 are renumbered generations that produced the two temporal samples. $t$ = gen2 – gen1 is elapsed time in generations between samples. $L$ = number of polymorphic loci used in each comparison; $F$ = Pollak's estimator of temporal $F$; $F$prime = $F$ adjusted for sampling error; Sprime is harmonic mean sample size across years and loci; $N_e$ is estimated $N_e$ using Equation 5.

| yr1 | yr2 | gen1 | gen2 | t | L | F | Fprime | Ne | Sprime |
|---|---|---|---|---|---|---|---|---|---|
| 2013 | 2014 | 1 | 2 | 1 | 988 | 0.02309 | 0.00245 | 204 | 48.45 |
| 2013 | 2015 | 1 | 3 | 2 | 959 | 0.02281 | 0.002 | 500 | 48.05 |
| 2013 | 2016 | 1 | 4 | 3 | 949 | 0.02449 | 0.00341 | 439 | 47.44 |
| 2013 | 2017 | 1 | 5 | 4 | 958 | 0.02439 | 0.00354 | 565 | 47.96 |
| 2013 | 2018 | 1 | 6 | 5 | 953 | 0.02133 | -0.00013 | -18806 | 46.6 |
| 2013 | 2019 | 1 | 7 | 6 | 965 | 0.02455 | 0.00357 | 840 | 47.66 |
| 2013 | 2020 | 1 | 8 | 7 | 906 | 0.02423 | 0.00271 | 1294 | 46.47 |
| 2014 | 2015 | 2 | 3 | 1 | 985 | 0.02173 | 0.00121 | 415 | 48.73 |
| 2014 | 2016 | 2 | 4 | 2 | 968 | 0.0236 | 0.00278 | 360 | 48.03 |
| 2014 | 2017 | 2 | 5 | 3 | 984 | 0.02412 | 0.00351 | 428 | 48.52 |
| 2014 | 2018 | 2 | 6 | 4 | 995 | 0.02366 | 0.00296 | 675 | 48.31 |
| 2014 | 2019 | 2 | 7 | 5 | 999 | 0.02433 | 0.0036 | 694 | 48.24 |
| 2014 | 2020 | 2 | 8 | 6 | 960 | 0.02553 | 0.00417 | 719 | 46.82 |
| 2015 | 2016 | 3 | 4 | 1 | 934 | 0.02222 | 0.00135 | 371 | 47.92 |
| 2015 | 2017 | 3 | 5 | 2 | 956 | 0.02126 | 6.00E-04 | 1673 | 48.4 |
| 2015 | 2018 | 3 | 6 | 3 | 964 | 0.0226 | 0.00175 | 857 | 47.96 |
| 2015 | 2019 | 3 | 7 | 4 | 962 | 0.02355 | 0.00269 | 744 | 47.94 |
| 2015 | 2020 | 3 | 8 | 5 | 924 | 0.0249 | 0.00349 | 717 | 46.71 |
| 2016 | 2017 | 4 | 5 | 1 | 924 | 0.02293 | 0.00203 | 247 | 47.85 |
| 2016 | 2018 | 4 | 6 | 2 | 946 | 0.02402 | 0.00286 | 350 | 47.26 |
| 2016 | 2019 | 4 | 7 | 3 | 937 | 0.02473 | 0.00359 | 418 | 47.3 |
| 2016 | 2020 | 4 | 8 | 4 | 896 | 0.02469 | 0.00297 | 673 | 46.04 |
| 2017 | 2018 | 5 | 6 | 1 | 956 | 0.0234 | 0.00245 | 204 | 47.73 |
| 2017 | 2019 | 5 | 7 | 2 | 955 | 0.02428 | 0.00331 | 302 | 47.69 |
| 2017 | 2020 | 5 | 8 | 3 | 920 | 0.02669 | 0.00512 | 293 | 46.36 |
| 2018 | 2019 | 6 | 7 | 1 | 961 | 0.02145 | 0.00048 | 1047 | 47.69 |
| 2018 | 2020 | 6 | 8 | 2 | 919 | 0.02179 | -0.00017 | -5868 | 45.54 |
| 2019 | 2020 | 7 | 8 | 1 | 905 | 0.02283 | 0.00136 | 367 | 46.58 |

Table S2. Simulated data used in the fits. Data for generations 2-4 were used to generate Equation 12; data for generations 1 and 7 (the Ends) were used to generate Equation 13. Ratio = $\sigma_{\hat{F}^{Adj}}/\sigma_{\hat{F}}$ is the ratio of the standard deviation of $\hat{F}^{Adj}$ to the standard deviation of the initial, single-generation $\hat{F}$.

Only first few lines are shown …

| Seed | Nreps | Generation | Ne | SampSize | Loci | Ratio |
|---|---|---|---|---|---|---|
| 4475 | 1000 | 1 | 50 | 25 | 100 | 0.863151 |
| 4475 | 1000 | 2 | 50 | 25 | 100 | 0.640535 |
| 4475 | 1000 | 3 | 50 | 25 | 100 | 0.650696 |
| 4475 | 1000 | 4 | 50 | 25 | 100 | 0.658716 |
| 4475 | 1000 | 5 | 25 | 25 | 100 | 0.686193 |
| 4475 | 1000 | 6 | 100 | 25 | 100 | 0.631432 |
| 4475 | 1000 | 7 | 50 | 25 | 100 | 0.824983 |
| 8486 | 1000 | 1 | 50 | 25 | 600 | 0.895657 |
| 8486 | 1000 | 2 | 50 | 25 | 600 | 0.635983 |

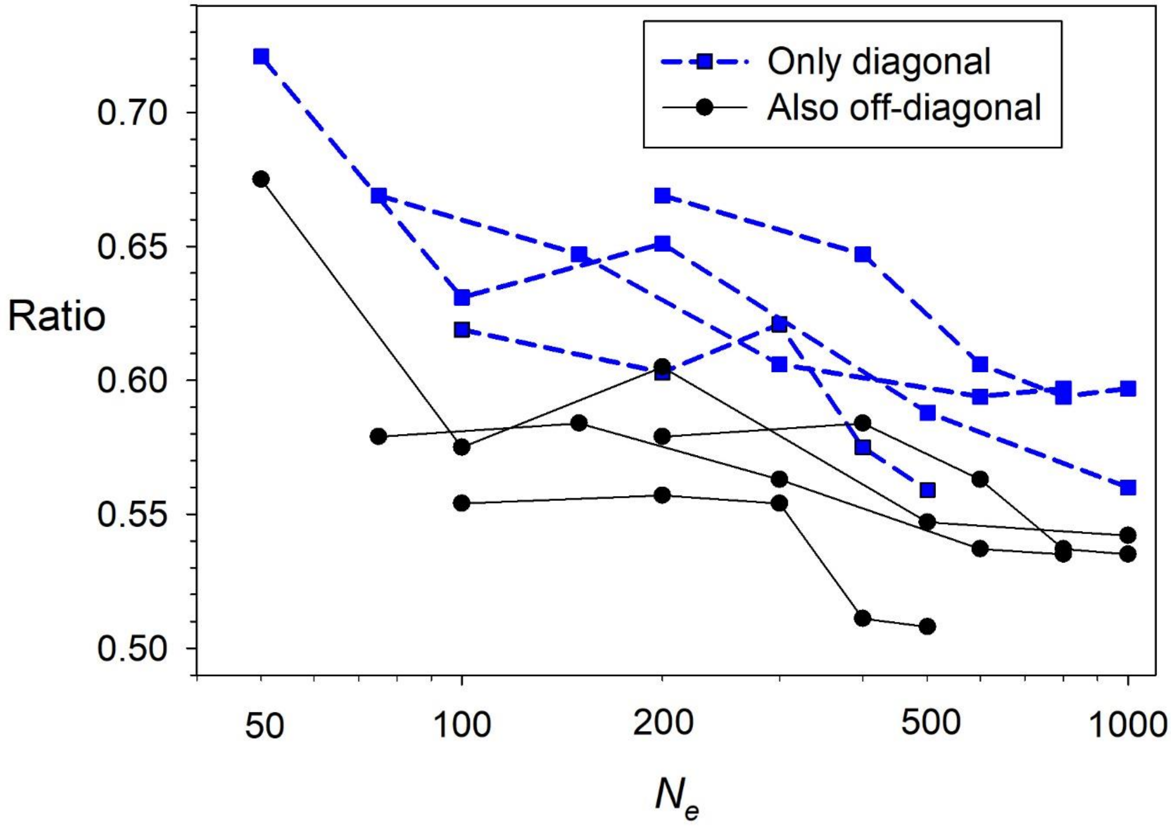


Figure S1. Simulation results evaluating benefits of iteratively updating off-diagonal elements (multigeneration $\hat{F}$) as well as single-generation (diagonal) estimates. Ratio = $\sigma_{\hat{F}^{Adj}}/\sigma_{\hat{F}}$ is the ratio of the standard deviation of $\hat{F}^{Adj}$ to the standard deviation of the initial, single-generation $\hat{F}$. Note the log scale on the X axis.

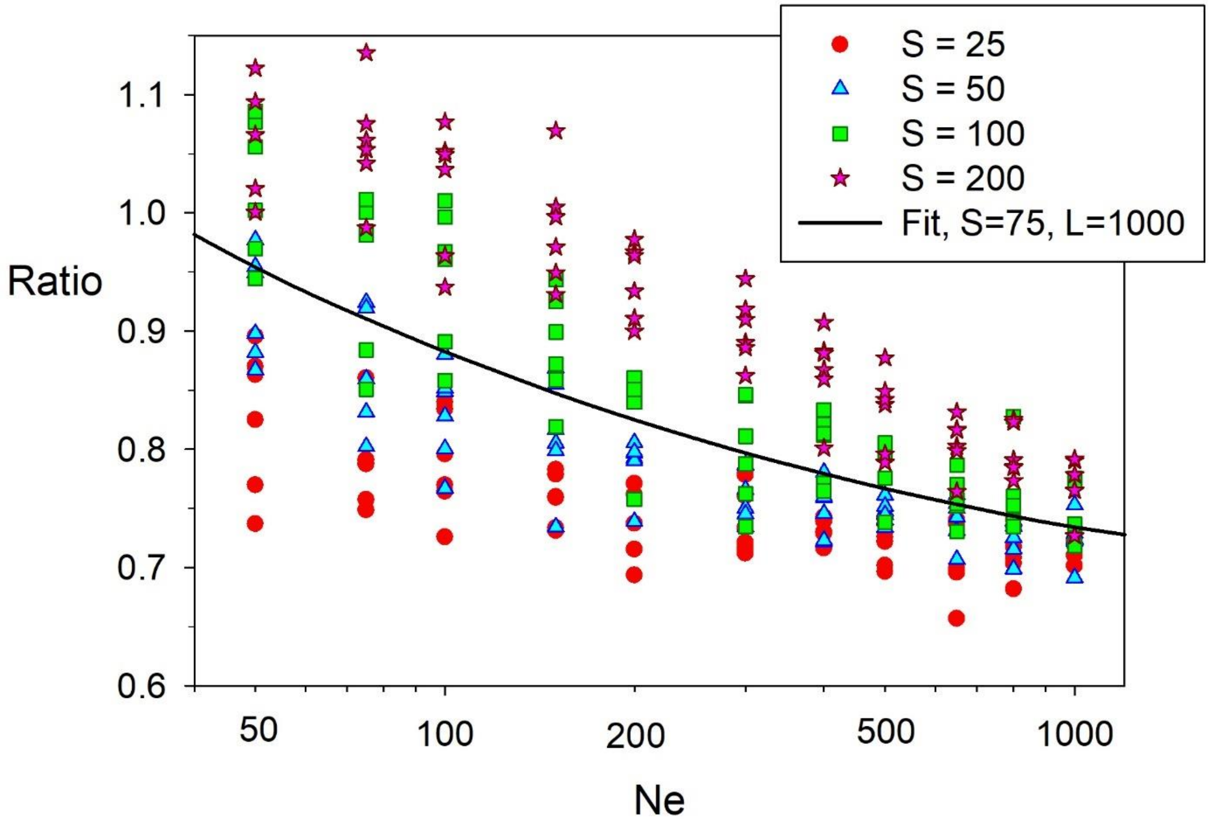


Figure S2. As in Figure 4 top, but for estimates that can leverage information from one additional generation of genetic drift either before or after the focal generation, but not both. Ratio = $\sigma_{\hat{F}^{Adj}}/\sigma_{\hat{F}}$ is the ratio of the standard deviation of $\hat{F}^{Adj}$ to the standard deviation of the initial, single-generation $\hat{F}$. Note the log scale on the X axis.
